# A Micro-scale Humanized Ventilator-on-a-Chip to Examine the Injurious Effects of Mechanical Ventilation

**DOI:** 10.1101/2024.02.26.582200

**Authors:** Basia Gabela-Zuniga, Vasudha C. Shukla, Christopher Bobba, Natalia Higuita-Castro, Heather M. Powell, Joshua A. Englert, Samir N. Ghadiali

## Abstract

Patients with compromised respiratory function frequently require mechanical ventilation to survive. Unfortunately, non-uniform ventilation of injured lungs generates complex mechanical forces that lead to ventilator induced lung injury (VILI). Although investigators have developed lung-on-a-chip systems to simulate normal respiration, modeling the complex mechanics of VILI as well as the subsequent recovery phase is a challenge. Here we present a novel humanized in vitro ventilator-on-a-chip (VOC) model of the lung microenvironment that simulates the different types of injurious forces generated in the lung during mechanical ventilation. We used transepithelial/endothelial electrical resistance (TEER) measurements to investigate how individual and simultaneous application of the different mechanical forces alters real-time changes in barrier integrity during and after injury. We find that compressive stress (i.e. barotrauma) does not significantly alter barrier integrity while over-distention (20% cyclic radial strain, volutrauma) results in decreased barrier integrity that quickly recovers upon removal of mechanical stress. Conversely, surface tension forces generated during airway reopening (atelectrauma), result in a rapid loss of barrier integrity with a delayed recovery relative to volutrauma. Simultaneous application of cyclic stretching (volutrauma) and airway reopening (atelectrauma), indicate that the surface tension forces associated with reopening fluid-occluded lung regions is the primary driver of barrier disruption. Thus, our novel VOC system can monitor the effects of different types of injurious forces on barrier disruption and recovery in real-time and can be used to identify the biomechanical mechanisms of VILI.

## Introduction

Mechanical ventilation (MV) is a critical form of life support for patients with respiratory failure and patients that require general anesthesia for surgical procedures.^1^ Patients with compromised respiratory function require MV to prevent hypoxemia, maintain acid-base balance, and alleviate the work of breathing associated with a pulmonary or systemic injury.^2^ While lifesaving, positive pressure MV can exacerbate existing lung injury and lead to ventilator induced lung injury (VILI).^3^ VILI arises from three primary biomechanical forces including alveolar over-distention (volutrauma), high transpulmonary pressure (barotrauma), and cyclic collapse/reopening of lung units (atelectrauma).^4^ Lung protective strategies that utilize low tidal volumes and modulate the positive end expiratory pressure (PEEP) have been employed to minimize VILI. However, patients requiring MV often have heterogenous lung injury, and non-uniform ventilation makes it difficult to eliminate the biomechanical forces that cause VILI.^5, 6^ Although animal models have been used to simulate VILI, these in vivo models are limited because they do not allow investigators to control and/or determine the relative importance of different biomechanical forces during VILI, cannot be used to monitor barrier function in real-time, and do not use human cells. Therefore, the development of novel in vitro models of VILI that can independently investigate the effect of the biomechanical forces that cause VILI, utilize primary human cells, and account for the clinical course of disease are needed.

Historically, in vitro models of VILI have utilized monoculture systems with a single cell type that allow for compressive stress^7, 8^ (barotrauma) or injurious strain via a flexible membrane^9, 10^ (volutrauma). Investigators have also modeled the surface tension and fluid shear stress (atelectrauma) that occur during airway/alveolar reopening by propagating an air liquid interface over an epithelium cultured on a static, rigid membrane.^10-12^ While these models provide valuable insight into injury patterns during MV, they have significant limitations including the lack of a fibrous extracellular matrix that mimics the basement membrane, lack of epithelial-endothelial interactions, and an inability to monitor barrier integrity in real-time. Furthermore, previous systems could not model multiple forms of VILI simultaneously and cannot monitor repair and recovery following injury.

Over the last several years investigators have developed microfluidic lung-on-a-chip (LOC) systems to try and recapitulate the alveolar microenvironment. Many of these models use co-cultures of transformed or neoplastic epithelial and endothelial cells that cyclically stretch a non-fibrous, homogenous, flexible membrane to mimic physiologic breathing.^9, 13^ These systems have been used to study lung cancer,^14, 15^ chronic obstructive pulmonary disease^16^ and asthma^17^. Ongoing advancements in micromachining technology have allowed investigators to use techniques like electrospinning to create nanofiber meshes that reflect the fibrous architecture of the native lung basement membrane.^18-20^ In parallel, other investigators have developed models of the alveolar microenvironment that mimic the alveolar geometry.^21, 22^ Although these advanced LOC models can simulate the physiologic structure and environment of the alveolus, none of them have been used to model the injurious physical forces that occur during mechanical ventilation. In addition, it is well known that tissue repair is a critical component of recovery following lung injury. However, previous LOC systems do not allow for investigation of how mechanical forces alter the degree of repair and/or the long-term recovery of barrier integrity. Therefore, the goal of this study was to develop a microfluidic ventilator-on-a-chip (VOC) model of the lung microenvironment that uses primary human cells, recapitulates the topography alveolar basement membrane, allows for the independent and concurrent simulation of injurious physical forces that occur during mechanical ventilation, and enables monitoring of barrier recovery during and following injury.

## Methods

### Fabrication & Assembly of the Ventilator-on-a-Chip

The device consists of two identical molds made of polydimethylsiloxane (PDMS; Sylgard 184, Dow Corning, Midland, MI) formed with a 10:1 base to curing agent ratio. As shown in Fig. 1A-C, the design features two concentric chambers separated by a 1 mm thick wall. The outer compartment is 5 mm wide and serves as the vacuum chamber which contains a single 18-gauge port that enables stretching of an electrospun membrane via negative pressure. The inner chamber, where cells are cultured, is 14 mm in diameter (Fig. 1C) and features two 18-gauge ports placed 2-3 mm from the chamber wall on opposite sides. The depth of both chambers, including the 1 mm thick inner chamber wall, is 4 mm unless otherwise stated. Platinum electrodes were incorporated into the device in a four-electrode configuration via direct insertion through the top of each PDMS mold. Wires were bent at a right angle so they could be easily secured to the top of their respective PDMS mold. During assembly, the electrospun membrane (see below) was attached to one half of the device and both halves were oxygen plasma treated. Both halves were then sealed together and allowed to cure, with additional PDMS, undisturbed for at least 2 hours before sterilization. Devices were sterilized by exposure to ultraviolet light for 10 minutes on each side. Devices were then flooded with 70% ethanol and left overnight. Prior to cell seeding, ethanol was removed, and devices were flooded with sterile PBS to ensure full hydration of the membrane overnight. For a subset of experiments simulating atelectrauma or combined volutrauma/atelectrauma, both sides of the electrospun membrane were collagen coated with type 1 bovine collagen (Advanced Biomatrix; Carlsbad, CA) overnight at 60 *µ*g/mL.

**Figure 1.**
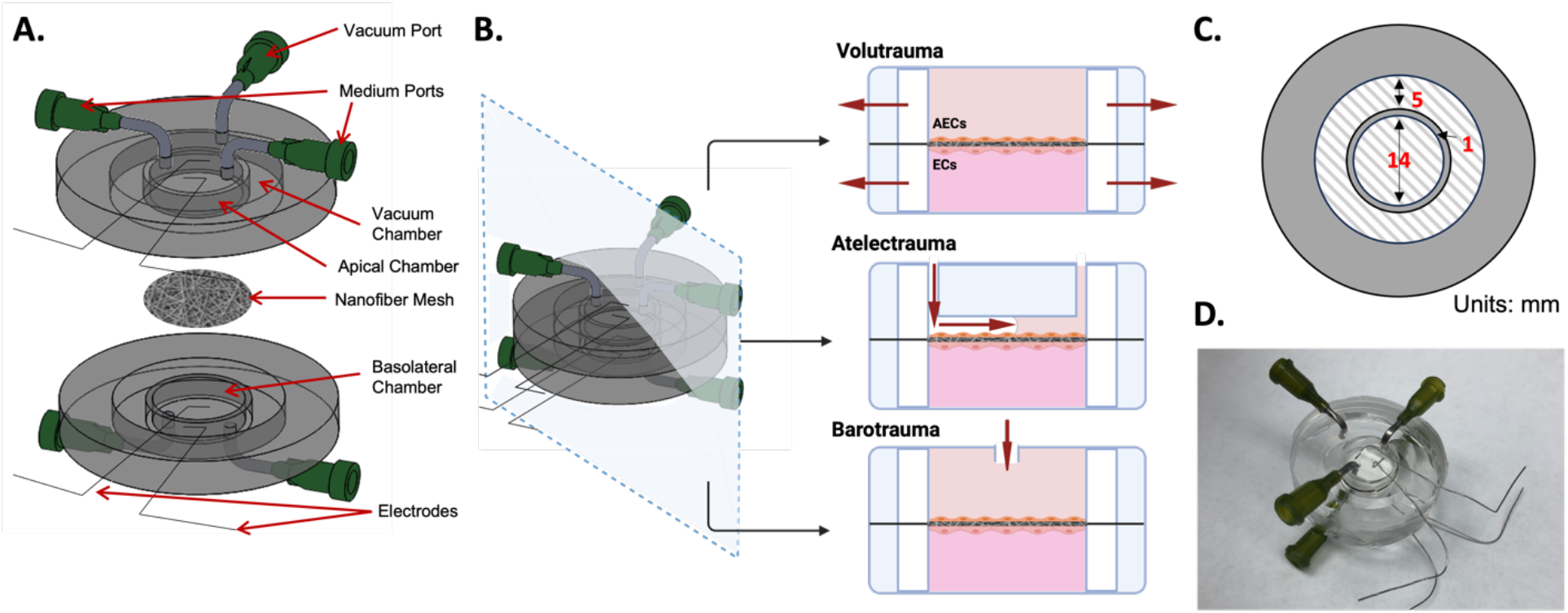
The ventilator-on-a-chip (VOC) design and working principle. **A**. Exploded view of VOC schematic. The device contains two concentric chambers; the vacuum compartment surrounds the inner circle where apical and basolateral chambers are separated by a nanofiber mesh. A four-electrode configuration of platinum wires is used to determine barrier formation via transepithelial/endothelial electrical resistance (TEER). **B**. Primary human lung epithelial and endothelial cells were co-cultured on opposite sides of the porous, electrospun polyurethane membrane. Volutrauma was simulated by applying injurious, cyclical, radial stretch. Atelectrauma was performed by propagating an air-liquid interface over the epithelium and barotrauma was achieved via cyclically pressurizing the apical chamber. Red arrows indicate the direction in which the force/injury was applied. **C**. Dimensions of described compartments. Top-down view. **D**. Image of VOC. AEC: alveolar epithelial cells, EC: endothelial cells.

### Electrospinning

A 5 weight/volume% polyurethane (PU; ChronoFlex C; AdvanSource; Wilmington, MA) solution was made by adding PU pellets to 1,1,1,3,3,3-Hexafluoro-2-propanol (HFP, Sigma-Aldrich; St. Louis, MO). A magnetic stir bar was used to mix the solution until the pellets were fully dissolved. A 10 mL syringe filled with the polymer solution and an 18-gauge flat tipped stainless-steel needle was placed in a syringe pump. The tip of the needle was 20 cm from the collector covered with aluminum foil (3” x 3”), which was connected to a high voltage generator while the needle tip was grounded. When 20 kV of positive voltage was applied, the polymer solution formed a Taylor cone that produced randomly aligned fibers (Fig. 2A). The syringe pump was programmed with a flow rate of 2.0 mL/hr. and scaffolds were spun for 24 minutes. Temperature and relative humidity were kept constant at 65-70° F and 20-25%, respectively.

**Figure 2.**
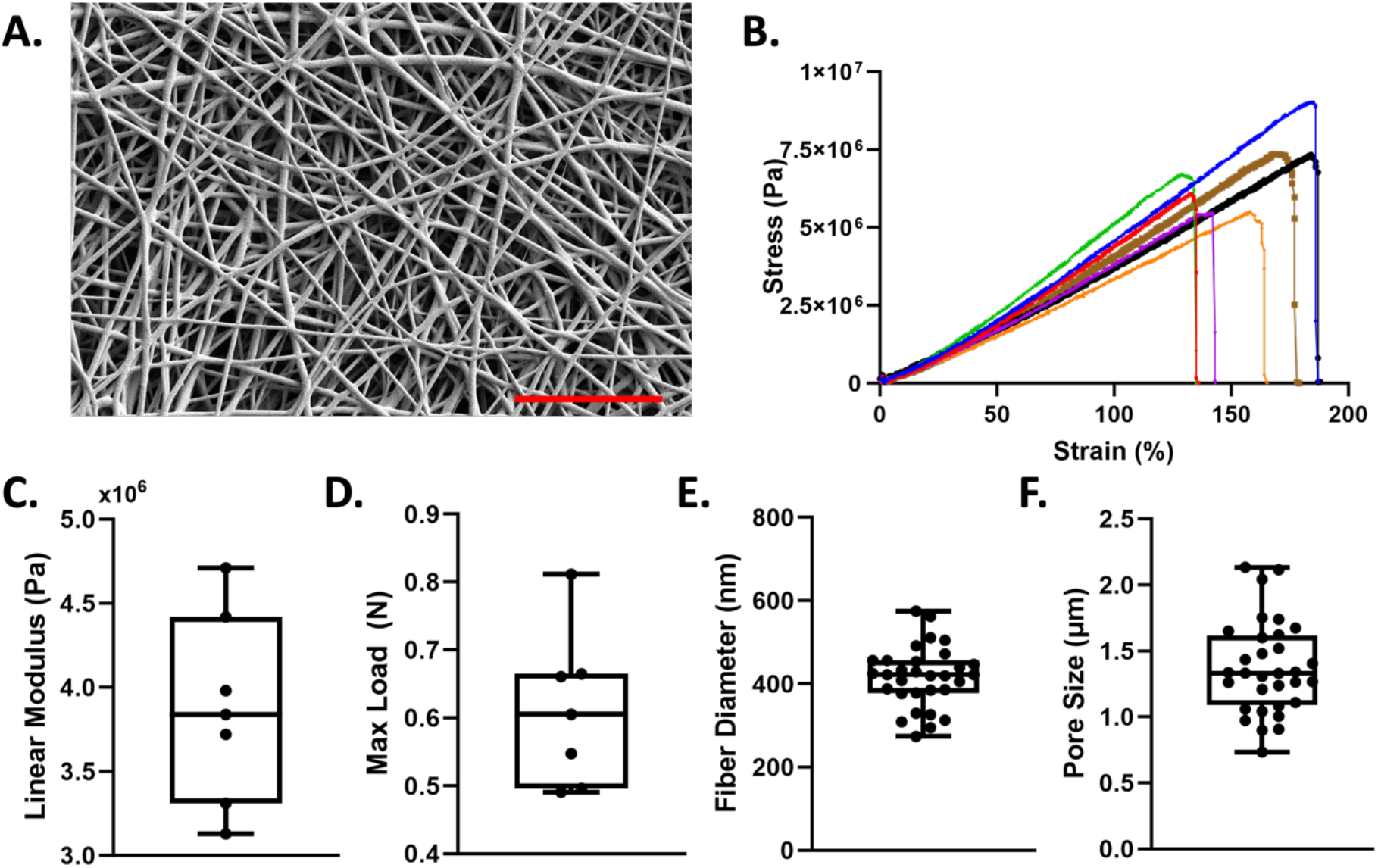
Nanofiber membrane fabrication and characterization. **A**. SEM image of 5 wt/vol% polyurethane membrane. Scale bar= 10 *µ*m **B**. Uniaxial tensile testing of n=7 polyurethane scaffolds illustrating the stress-strain curves. Quantification of the **C**. linear modulus and **D**. maximum load. ImageJ was used to quantify **E**. fiber diameter and **F**. pore size. Min to max all points shown.

### Transepithelial/endothelial electrical resistance (TEER) measurements

A Vertex.One.EIS potentiostat (Ivium Technologies, the Netherlands) was used to record impedance spectra. Four platinum electrodes were used to obtain four-point impedance measurements by applying a 1 Hz, 10 *µ*A AC current to the current leads and measuring voltage in real-time via the voltage leads. Impedance was calculated by 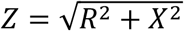 where Z is the magnitude of the impedance in ohms, R is the real part and X is the imaginary part of the impedance. Low frequency values were reported because TEER dominates the signal at lower frequencies (<100 Hz)^23^. Data are also presented as a percent change of impedance (%ΔZ), where values obtained at each time point were normalized to the 0 hr. timepoint.

### Membrane Characterization

Morphology of the electrospun fibers was examined using scanning electron microscopy (SEM). The Apreo SEM was operated at an accelerating voltage of 2 kV and current of 50 pA. The 2D pore size and fiber diameter were quantified in ImageJ using three samples (n=1 per sample) and 30-40 measurements per image. Pore size was assessed by measuring the interfiber distance, e,g. the distance between opposing sides of two fibers that formed a single 2D pore. Membrane thickness was characterized using a Helios focused ion beam (FIB) laser. The laser was used to mill into the fibers to create a trench that exposed a clean interface between the fiber mesh and the underlying aluminum foil. The SEM was then used to measure the thickness of the fiber scaffold. Three measurements from three random regions were selected and measured.

### Tensile Testing

The mechanical properties of the electrospun fibers membrane were quantified via tensile testing. A dog-bone shaped punch, 3 mm width and 27 mm length, was used for all samples. Samples were hydrated in PBS prior to strain tests. Samples were strained at a grip speed of 2 mm/sec until failure using a tensile tester (TestResources 100R; Shakopee, MN). Stress-strain curves were generated for n=7 samples across two different scaffolds. Cross-sectional area of the membrane and the slope of the stress-strain curve in the 50-70% strain region (i.e. linear region) was used to determine the linear elastic modulus.

### Cell-Culture

Primary human alveolar epithelial cells (pneumocytes) were obtained from Cell Biologics and cultured in human epithelial cell (HEC) growth media supplemented with a human epithelial supplement kit. Primary human lung microvascular endothelial cells (HMVEC-L) were obtained from Lonza and cultured in endothelial cell basal medium (EBM-2) and supplemented with a microvascular endothelial cell growth media (EGM2-MV) kit. All cell types and donors were non-smokers and grown according to manufacturer’s recommendations. Prior to cell seeding in the VOC, ‘Day 0’ TEER measurements were obtained to calibrate baseline impedance of the nanofiber membrane. Once cells were grown to confluence on tissue culture plastic, pneumocytes were collected and injected into the apical chamber at a density of 3.5 x 10^5^ cells/mL. Cells were allowed to adhere under static conditions for 4-5 hours or overnight. The device was then inverted, and the HMVEC-Ls were injected into the basolateral chamber at a density of 6x10^4^ cells/mL and allowed to adhere overnight under static conditions. Cells were grown under a submerged cultured at 37°C and media was manually changed every other day unless otherwise stated. Media was allowed to equilibrate for 15-20 minutes at 37°C prior to taking TEER measurements. Barrier formation was assessed via TEER measurements taken every other day (during media changes) until impedance measurements reached a plateau (+/-100 Ω) over 2-4 days indicating the formation of a complete barrier.

### Simulation of Ventilator-Induced Lung Injury

Prior to all experiments, fresh media was introduced to both apical and basolateral chambers and allowed to equilibrate for 15-20 minutes at 37°C. All experiments were completed at 37°C.

#### Atelectrauma

Atelectrauma was simulated by propagating an air-liquid interface over the epithelial layer. Since previous investigators^24^ have demonstrated that atelectrauma occurs in the distal small airways and alveoli with diameters <2 mm, we modified the apical chamber geometry (Fig. 1) so that the height of the apical chamber in which airway reopening occurs was approximately 2 mm. All devices used for atelectrauma experiments reflect this change in geometry. For these studies, the apical chamber was perfused at a flow rate of 0.05 mL/min every other day using a syringe pump and the basolateral chamber media was changed manually as described. Atelectrauma was simulated using previously described techniques.^18^ Briefly, a programmable Harvard PhD 2000 syringe pump was used to first retract fluid at either 10.5 mL/min and 5.25 mL/min to create a “forward”-propagating air-liquid bubble at a velocity of 5 mm/sec or 2.5 mm/sec respectively. After reopening, the channel was then refilled with fluid at the same respective rate and the cycle was repeated for 2 hours. Note the above flow rates/velocities correspond to respiration frequencies of 0.25 Hz and 0.125 Hz, respectively. The two selected frequencies are based on normal tidal volume ventilation and previous work that demonstrates slower frequencies result in more injury^25^. Note, these reopening velocities correspond to the expected reopening velocities of 1-10 mm/s for normal breathing conditions (V_T_= 500 mL, 12-15 breaths/min) and total cross-sectional area for terminal/respiratory bronchioles (100-1000 cm^2^).^26^ TEER measurements were taken periodically, at 30, 60, 90 and 120 minutes to monitor barrier function. Following injury, recovery was monitored for 48 hours. Separate experiments were conducted to assess cell death (i.e. plasma membrane rupture) in response to reopening velocity/frequency. For these experiments, atelectrauma was simulated for 2 hours and a live/dead cell viability assay and ImageJ was used to quantify the percentage of live and dead cells.

#### Barotrauma

Transmural pressure was simulated by applying oscillatory air pressure to the apical chamber via a central port (Fig. 1). The central port was connected to a water manometer and a small animal ventilator (Harvard Apparatus, Holliston, MA). Consistent with previous studies from our group^8^, oscillatory pressure was applied for 24 hours with a triangle waveform from 0-20 cm H_2_O at 0.2 Hz at 37°C. TEER measurements were taken periodically for 6 hours and then again at 24 hours. During these experiments, all other ports were closed including the vacuum chamber so that no pressure can be dissipated through the membrane or basolateral chamber.

#### Volutrauma

To simulate volutrauma, cells were cyclically stretched using a computer controlled vacuum pump (FlexCell International; Burlington, NC) and subjected to two either 10% or 20% radial strain at a frequency of 0.25 Hz for 4 hours at 37°C. These strains are equivalent to 21% and 44% area strain (i.e. change in area/initial area) and thus represent strains expected at large lung volumes (i.e. ∼80% and >100% total lung capacity)^27^. During volutrauma experiments, TEER measurements were taken periodically for 4 hrs. and recovery measurements were assessed for 48 hours. Membranes were calibrated to determine how much pressure was needed to generate desired strain levels as shown in supplementary Fig. 1. Six devices were used for calibration purposes and for these studies, fluorescent beads (diameter= 2 *µ*m) diluted in PBS were flooded into the apical chamber and beads were allowed to attach to the fibrous membrane overnight. A syringe pump was used to withdraw air from the vacuum chamber at 1 mL/min which generated a negative vacuum pressure that linearly decreased from 0 to -7 PSI. Prior to membrane stretching, each device was secured to the stage of an inverted epifluorescent microscope, and the displacement of beads located 1, 3, 5, and 7 mm from the center of the membrane was measured using ImageJ. Radial strain was calculated by dividing the displacement by the original radial position.

#### Simultaneous Volutrauma and Atelectrauma

Volutrauma and atelectrauma were simulated simultaneously as described above with a 2 mm channel height under two different conditions. Volutrauma was simulated by applying 10% cyclic radial strain via an appropriate vacuum pressure and simultaneously atelectrauma was simulated by propagating and air-liquid interface over the epithelium at 10.5 mL/min flow rate (0.25 Hz) or 5.25 mL/min (0.125 Hz) for 2 hours. Note that both volutrauma and atelectrauma were synchronized where cyclic strain was performed at the same frequency of atelectrauma with no phase offset. TEER measurements were taken periodically to assess barrier function and following injury, recovery was monitored for 48 hours.

### Immunofluorescence Microscopy

To visualize tight junction formation, cells were fixed in 4% PFA for 10 min and permeabilized with 0.1% Triton X-100 in PBS for 10 min. Cells were blocked with 1% BSA/5% Goat Serum solution for 1 hour at room temperature and then stained with anti-ZO-1 antibody (1:200; Invitrogen; Carlsbad, CA) for pneumocytes and anti-VE-cadherin (1:100; Invitrogen; Carlsbad, CA) for HMVEC-Ls overnight at 4°C. The following day, an Alexa Fluor 488 conjugated secondary antibody (1:500, goat anti-rabbit; Thermo Fischer, Waltham, MA) was added for 1 hour and cells were counterstained with Hoechst (1:10,000). The membrane was then removed from the device and mounted to a glass coverslip. Images were taken with a Leica Stellaris 5 confocal microscope at 120X. To visualize monolayer formation for both epithelial and endothelial cells, a Phalloidin stain was employed and counterstained with Hoechst. Cells were fixed in 4% PFA, permeabilized with Triton X-100, blocked with 1% BSA/5% goat serum and stained at 1:40 Phalloidin for 1 hour. Cells were then counterstained with Hoechst (1:10,000) and the membrane was mounted to a glass coverslip. Images were taken with a Leica Stellaris 5 confocal microscope at 60X.

### Statistics

Statistical analysis was performed using GraphPad Prism 10. All statistical analyses were performed on TEER measurements normalized to zero hr. timepoint measurements. Outliers were identified using the ROUT method with a 1% threshold.^8^ Data were tested for normality using a Shapiro-Wilk test. For data that were normally distributed, a one-way ANOVA was performed to compare multiple groups. For experiments with two independent variables, a two-way ANOVA was performed. If a significant effect was found, a Tukey or Dunnett’s post-hoc test for group comparisons was performed. A p-value of ≤< 0.05 was considered statistically significant. Error bars for all data presented in line and bar graphs is the standard deviation. Error bars for data presented as a box and whisker plot are min to max.

## Results

### Electrospun polyurethane scaffold characterization

Medical grade, thermoplastic polyurethane was chosen for its biocompatibility and deformablity^28^ and electrospun to create fibrous scaffolds (Fig. 2A). Scaffold characterization via uniaxial tensile testing (Fig. 2B) reveals polyurethane membranes have an average modulus of 3.8 MPa ± 0.11 (Fig. 2C) and can withstand an average maximum load of 0.6 N ± 0.56 (Fig. 2D). Average fiber diameter was 420 nm ± 99.43 (Fig. 2E) and average 2D pore size was 1.35 *µ*m ± 436 nm. (Fig. 2F), which is similar to lung basement membrane^29, 30^.

### The ventilator-on-a-chip (VOC) recapitulates the function of the alveolar-capillary barrier

Barrier formation was assessed by immunostaining for ZO-1 and VE-cadherin in epithelial and endothelial cells respectively. Confocal images in Fig. 3A and 3B indicate the presence of tight junctions and adherens junctions following 14 days of primary cell co-culture in the VOC. To visualize monolayer formation for both cell types, a z-stack image was obtained after staining both epithelial and endothelial cells with Phalloidin to label actin (Fig. 3C). These images demonstrate monolayer formation and separation between the apical and basolateral cell monolayers from the electrospun membrane. To assess barrier function we measured electrical impedance^23^ by obtaining TEER measurements every 2 days for 14 days in culture (Fig. 3D). Under co-culture conditions, TEER measurements began increasing at day 6-8 and reached >2000 Ω after 10 days. Interestingly, when only epithelial cells were cultured in the device, TEER measurements only increased to ∼1000 Ω by 6-8 days which was significantly lower than co-culture conditions and when only endothelial cells were cultured in the device, we observed no increase in TEER measurements. These data indicate that although epithelial and endothelial cells form monolayers (Fig 3A and 3B), cell-cell communication under co-culture conditions is required to obtain an electrically tight barrier in our VOC device.

**Figure 3.**
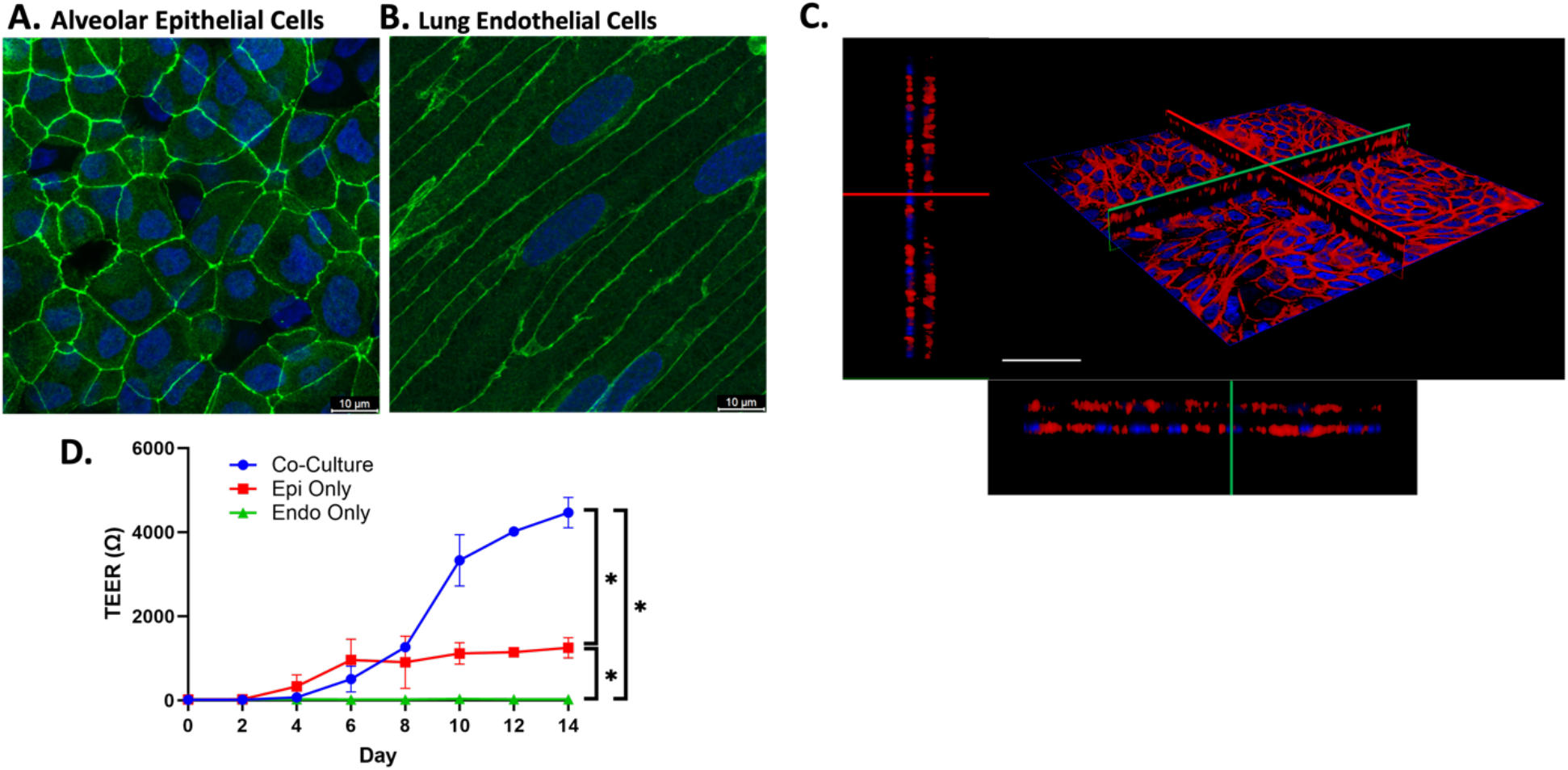
VOC recapitulates lung barrier function. Culture of alveolar epithelial cells and human lung microvascular endothelial cells in the VOC reveals confluent monolayers with formation of tight junctions as indicated by staining for **A**. ZO-1 in epithelial cells and adherens junctions in endothelial cells by staining for **B**. VE-Cadherin. **C**. Confocal image of co-cultured cells stained with Phalloidin (red) and Hoescht (blue) indicates that epithelial and endothelial cells are separated by the nanofiber membrane. Red and green lines indicate the X (upper left)-Y(lower right) intersection. Scale bar= 50 *µ*m. **D**. Barrier function was assessed by transepithelial electrical resistance (TEER) for co-culture and monoculture conditions. Data were analyzed via repeated measures two-way ANOVA with Tukey’s multiple comparison test. *p<0.0001

### Volutrauma leads to magnitude dependent barrier disruption and subsequent recovery

We used the VOC to investigate how different cyclic radial strain magnitudes alter both barrier disruption (reduction in TEER) and recovery (increase in TEER after injury). As shown in Fig. 4A, 20% radial strain results in a statistically significant drop in impedance relative to the 0 hr. timepoint after 30 minutes and TEER measurements remain low throughout 4 hours of injury. Stopping the injurious 20% radial strain resulted in an increased impedance value that was not statistically different from the zero-hour timepoint indicating improved barrier function within one hour. At 10% radial strain, we observed a transient decrease in barrier function following stretch (Fig. 4A) at 30 minutes and a recovery to baseline during the injury period. Static controls resulted in non-significant fluctuations in TEER measurements. Comparisons between different strain magnitudes (Fig. 4C) reveal a significant difference between all groups after 0.5 hours of injury with 20% radial strain exhibiting more injury than 10% radial strain. Following four hours of stretch, barrier integrity is significantly lower at 20% radial strain compared to 10% radial strain while the injury level at 10% radial strain is not different than static controls. By 5 and 24 hours (1 and 23 hours of recovery) there is no statistically significant difference in injury among the groups indicating that the alveolar-capillary barrier function rapidly recovers following volutrauma at all strain magnitudes.

**Figure 4.**
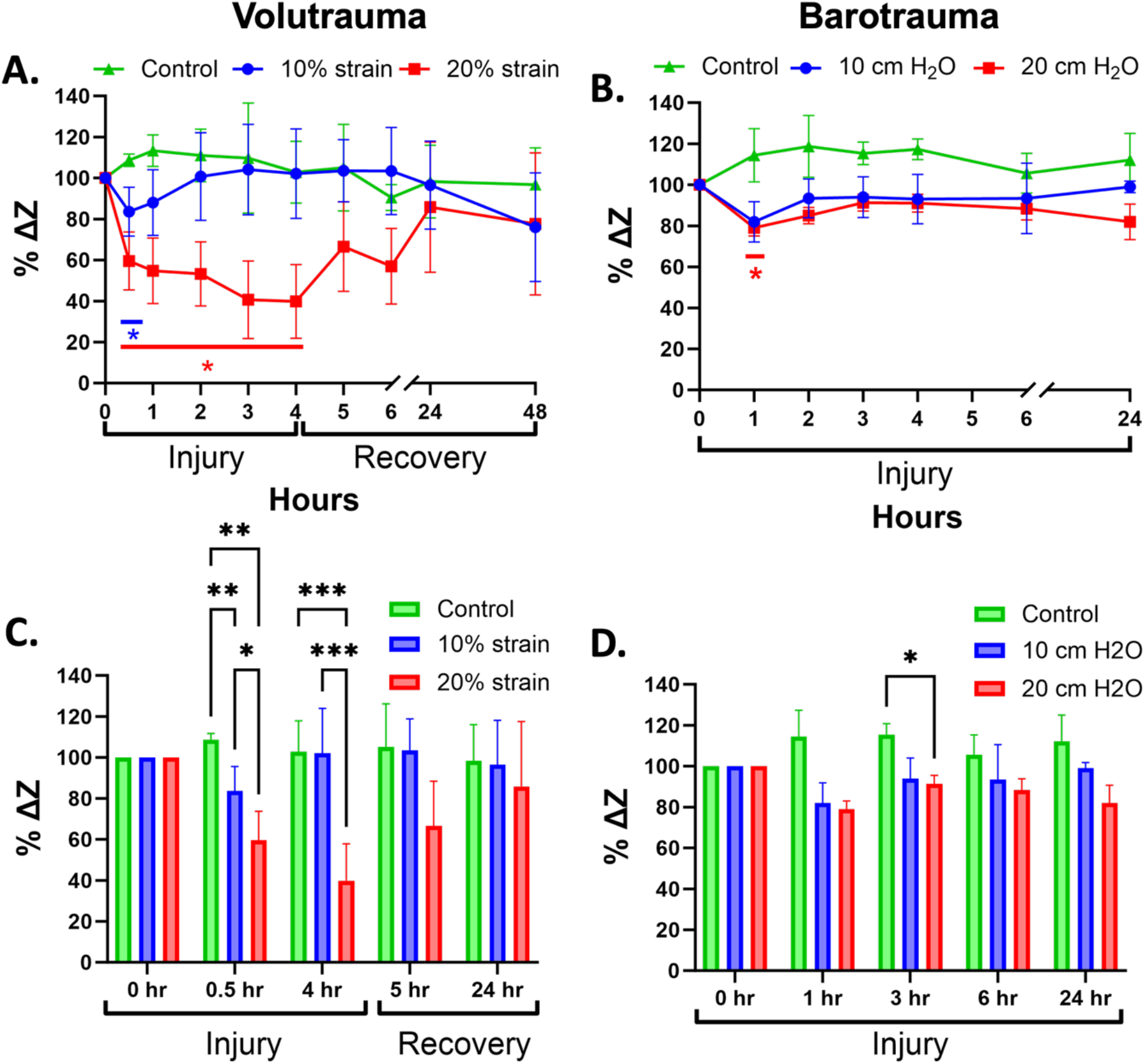
Volutrauma and barotrauma have a variable impact on barrier function. Barrier integrity as assessed by percent change of impedance (% ΔZ) for **A**. 4 hrs. of volutrauma and 48 hrs. of recovery for static controls (n=6), 10% strain (n=8) and 20% strain (n=5). Data are normalized to 0 hr. time point. Statistical differences for 20% and 10% radial strain are relative to their respective 0 hr. time points denoted as *p<0.05. Barrier integrity was also assessed for **B**. 24 hrs. of barotrauma for 10 cm H_2_O (n=3), 20 cm H_2_O (n=3) and static controls (n=3). Statistical differences for 20 cm H_2_O are relative to 0 hr. time point denoted as *p<0.05. All data for barotrauma and volutrauma are normally distributed by the Shapiro-Wilk test and analyzed by repeated measures one-way ANOVA with Dunnett’s multiple comparison test. Additional multiple comparisons via Tukey’s test were completed following a repeated measures mixed effects analysis (2-way ANOVA) for **C**. volutrauma and **D**. barotrauma. *p<0.05, **p<0.005, ***p<0.001. All data presented are shown as means with standard deviation.

### Barotrauma does not cause significant barrier disruption

As shown in Fig. 4B, we used the VOC to investigate how different oscillatory transmural pressure magnitudes alter barrier disruption. Application of high cyclic transmural pressure (20 cm H_2_O) resulted in a small but statistically significant decrease in relative impedance at the 1 hr. time point compared to the zero hour timepoint. However, the relative impedance at the 3, 6 and 24 hr. timepoints were not statistically different than the zero timepoint. In addition, 10 cm H_2_O resulted in no significant differences in relative impedance over 24 hours of injury. Comparisons between the different levels of oscillatory transmural pressure only indicated a significant difference between 20 cm H_2_O and static controls following three hours of cyclical pressure (Fig. 4D). These data indicate that this degree of barotrauma is less injurious than volutrauma in the VOC system.

### Atelectrauma leads to loss of barrier integrity with a delayed recovery compared to volutrauma

We used the VOC to investigate how different reopening velocities/frequencies during atelectrauma alter both barrier disruption and recovery. As shown in Fig. 5A, atelectrauma induces a very large and statistically significant reduction in relative impedance relative to the zero-hour timepoint within 30 minutes and this rapid and dramatic loss of barrier function is sustained during two hours of atelectrauma for both high and low frequencies. Interesting, after the cessation of atelectrauma, the relative impedance remains low where the impedance at the 3 and 4 hour timepoints remain statistically lower than the zero-hour impedance. Only after 22 to 46 hours of recovery (24 and 48 hr. timepoints) do we observe an improvement in barrier integrity and relative impedances that are not statistically different than the zero-hour timepoint. Comparisons between groups (Fig. 5B) reveal that at all injury time points and after one hour of recovery (3 hr. timepoint), both reopening frequencies resulted in significantly more barrier disruption compared to control conditions. However, the relative impedance measured at 0.125 Hz and 0.25 Hz during injury and recovery was not statistically different. After 48 hours of recovery, there were no statistically significant differences between any group. Since previous investigators have demonstrated that atelectrauma causes significant cell injury/plasma membrane rupture,^11, 12^ we used a Live/Dead assay to investigate how atelectrauma in the VOC influenced cell injury/plasma membrane rupture. As shown in Fig. 5C, we found significantly increased cell death at both frequency conditions with the slower frequency (0.125 Hz) resulting in significantly more plasma membrane rupture/cell death than the static control and faster frequency (0.25 Hz). Representative live/dead images support this quantitative data (Fig. 5D). These data indicate that in our VOC model, atelectrauma rapidly disrupts the barrier and causes cell death which delays repair/recovery after this form of VILI.

**Figure 5.**
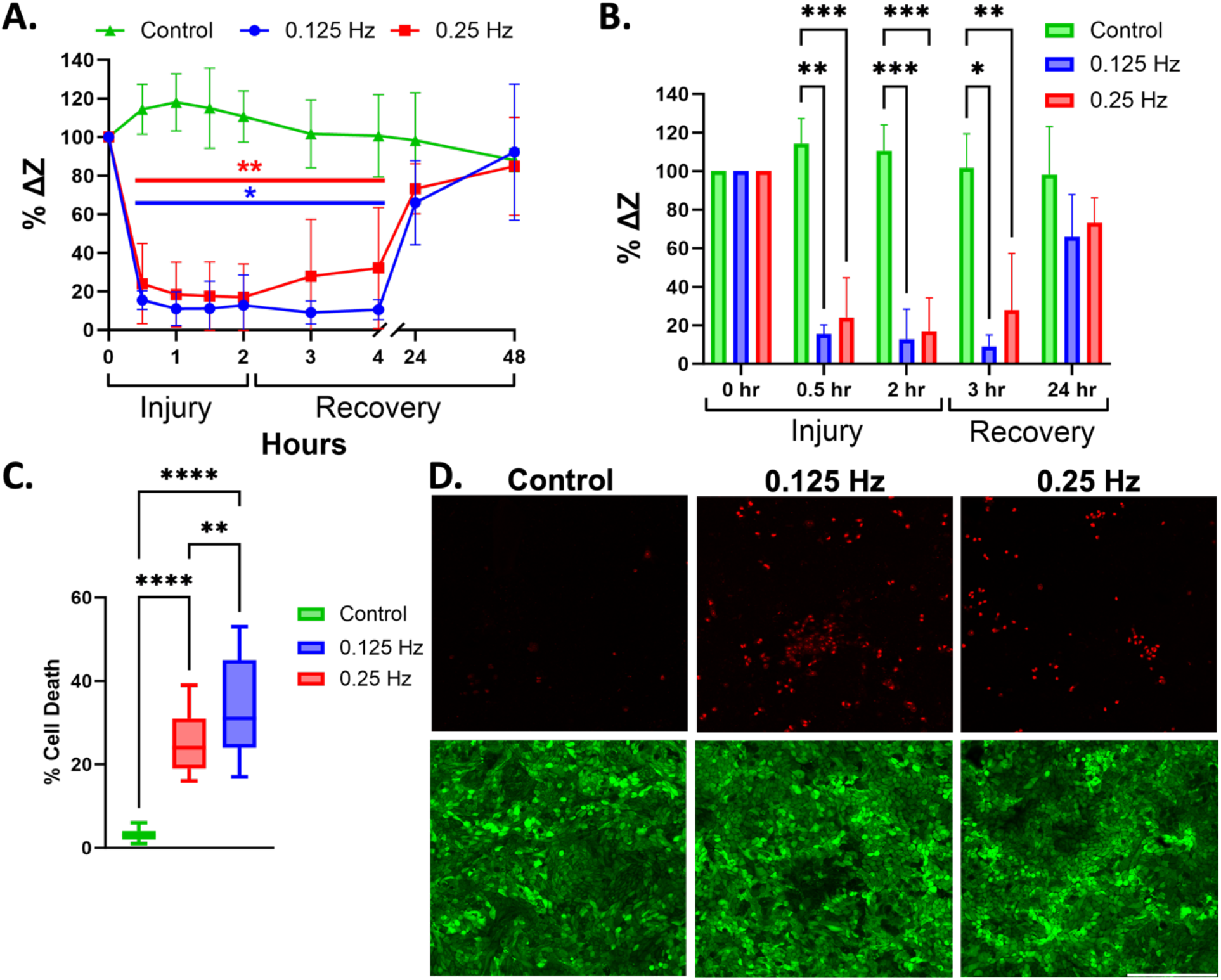
Atelectrauma leads to loss of barrier function and cell death. **A**. Barrier integrity as assessed by percent change in impedance (% ΔZ) for 2 hrs. of atelectrauma and 48 hrs. of recovery for static controls (n=3), 0.125 Hz (n=6) and 0.25 Hz (n=6). Data were normalized to 0 hr. time point. Data for each group were analyzed by repeated measures one-way ANOVA with Dunnett’s multiple comparisons test. Statistical differences for each frequency are relative to 0 hr. time point denoted as *p<0.05 and **p<0.01. **B**. Additional multiple comparisons via Tukey’s test were completed following a repeated measures two-way ANOVA. *p<0.05, **p<0.005, ***p<0.001. **C**. Devices were injured for 2 hours (n=2 per group) and quantified for percentage of cell death via manual cell counting in ImageJ. Data are normally distributed by Shapiro-Wilk test. Data were analyzed via repeated measures one-way ANOVA with Tukey’s multiple comparison test. Data is presented as min to max. ****p<0.0001 and **p<0.01. **D**. Representative Calcein/propidium iodide images at each condition (red=dead cells, green=live cells).Scale bar= 200 *µ*m. Data presented as means with standard deviation.

### Barrier disruption during simultaneous volutrauma and atelectrauma is driven by atelectrauma

Since patients undergoing MV are exposed to multiple injurious physical forces concurrently, we used the VOC to investigate how simultaneous application of volutrauma and atelectrauma impact barrier disruption. First, we simultaneously applied 10% radial strain and atelectrauma at 0.25 Hz in our VOC. This resulted in a rapid and statistically significant decrease in barrier function at 30 minutes which remained statistically lower than the zero-hour timepoint during the 2 hours of injury (Fig. 6A). In addition, after cessation of simultaneous volutrauma/atelectrauma, the relative impedance remained statistically lower than the zero-hour impedance for an additional 2 hours. Impedance was not statistically different than the zero-hour timepoint only after 24-48 hours of recovery. Although similar trends were observed when we only applied atelectrauma in the VOC (circles in Fig. 6A), volutrauma only (triangle in Fig. 6A) resulted in a different pattern. Specifically, volutrauma alone led to a small but statistically significant decrease in relative impedance at the 0.5 and 1-hour injury timepoints. However, impedance values at 1.5 and 2 hours were not statistically different than the zero-hour timepoint and recovery was rapid with all impedance values during recovery not statistically different than the zero-hour timepoint. We then repeated these experiments at 0.125 Hz and observed similar trends (Fig. 6B). One important difference in the lower frequency experiments was that volutrauma did not induce a significant reduction in impedance at any injury or recovery time point when compared to the zero-hour control and both atelectrauma and combined conditions resulted in more barrier disruption as evidence by relative impedance values <10%. Note that the data presented for atelectrauma only in Fig 6 is the same data from Fig 5.

**Figure 6.**
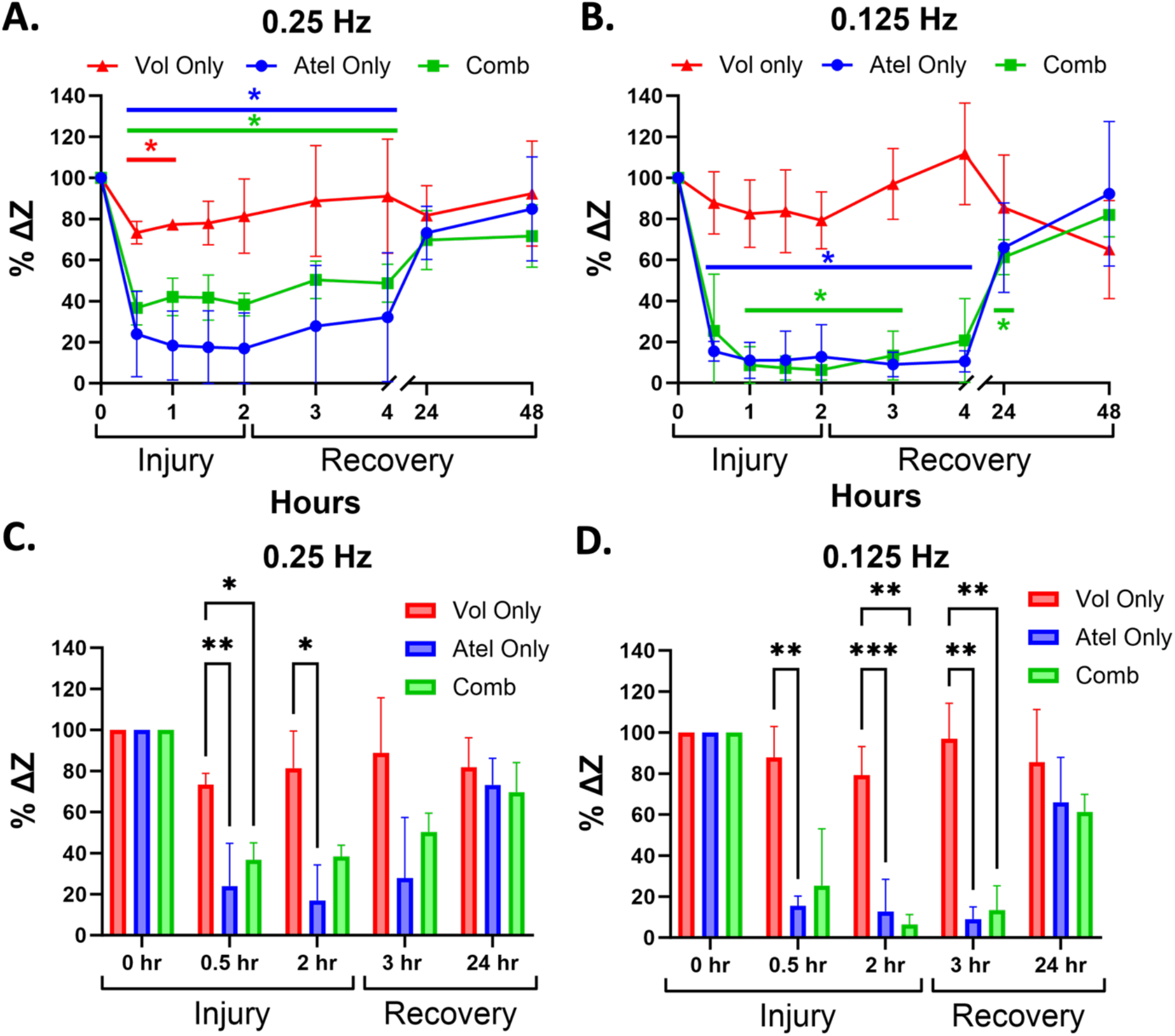
Injury frequency leads to variable impact on barrier function under combined injury conditions. Barrier integrity as assessed by percent change of impedance (% ΔZ) at **A**. 0.25 and **B**. 0.125 Hz during 2 hrs. of injury and 48 hrs. of recovery for volutrauma only (10% strain at 2 mm channel height; n=4), atelectrauma only (data from Fig. 4A; n=5) and combined volutrauma & atelectrauma (n=3). All data are normally distributed by Shapiro-Wilk test. Data were analyzed via repeated measures one-way ANOVA with Dunnett’s multiple comparison test. Data are normalized to 0 hr. time point. *p<0.05. Additional multiple comparisons via Tukey’s test were completed following a repeated measures two-way ANOVA for **C**. 0.25 Hz and **D**. 0.125 Hz. *p<0.05, **p<0.005, ***p<0.001. All data presented as means with standard deviation. Comb-combined volutrauma and atelectrauma; Vol only-volutrauma only; Atel only-atelectrauma only.

When comparing groups at the higher 0.25 Hz frequency, Fig. 6C reveals that atelectrauma and the combined conditions cause significantly more barrier disruption (lower impedance) than volutrauma at the 0.5-hour timepoint and that atelectrauma induced significantly more barrier disruption than volutrauma at 2 hours. We did not observe any significant differences in relative impedance during recovery between conditions at 0.25 Hz. At a slower frequency (0.125 Hz), simultaneous injuries and atelectrauma alone both resulted in significantly more barrier disruption than volutrauma after 2 hours of injury (Fig 6D) and during the initial recovery phase (3 hrs.). Importantly, there were no statistically significant differences between the atelectrauma and combined conditions at any injury or recovery timepoint. Together, this data indicates that atelectrauma is the driving physical force causing barrier disruption and that slower reopening of fluid occluded lung units may be the most important physical force to mitigate when trying to reduce the degree of VILI.

## Discussion

In this study, we developed a novel ventilator-on-a-chip (VOC) system that was specifically designed to investigate the complex biomechanics associated with ventilation induced lung injury (VILI). The VOC recapitulates the functional in vivo barrier properties of the alveolar capillary barrier including tight junction formation in epithelial cells, adherens junction formation in endothelial cells (Fig. 3A and 3B), monolayer formation (Fig. 3C), and the development of an electrically tight barrier (Fig. 3D). The electrospun nanofiber membrane used in the VOC exhibits several characteristics of the basal lamina, including pore size (Fig. 2F) and a fiber diameter (Fig. 2E) that are consistent with measurements of human collagen fibrils^29, 30^. Importantly, the VOC utilizes primary human epithelial and endothelial cells and exposes these cells to different types of physical forces that occur during mechanical ventilation of injured lungs. To our knowledge, our group is the first to develop an in vitro model of lung injury that accounts for primary cell-based co-culture, the fibrous texture of the lung basement membrane, and exposure of cells to the individual or concurrent physical forces responsible for VILI.

Advancements in micromachining technology have allowed investigators to develop sophisticated organ-on-a-chip systems to study a wide array of diseases. Huh et al. was the first to develop a lung-on-a-chip (LOC) platform with the capability to generate physiologic levels of uniaxial strain (5-15%) using primary endothelial cells and transformed lung epithelial cells cultured on a thin PDMS membrane.^31^ Several investigators have developed enhanced LOC models which can replicate negative pressure diaphragmatic breathing^13, 32^, mimic the in-vivo 3D geometry of alveolar sacs^22^ and incorporate nanofiber membranes to study transepithelial molecular transport^19^. However, none of these previous systems were designed to investigate how the complex biophysical forces generated during MV cause lung injury and although previous systems used TEER measurements to monitor barrier function,^23^ previous systems could not assess real-time changes in barrier function due to biomechanical injury, especially in the context of VILI. Therefore, the VOC model presented here represents a significant advance since it allows for the real-time assessment of how the biophysical forces associated with VILI alter barrier disruption.

The VOC presented here accounts for several of the key components required to form a functional alveolar-capillary barrier including primary cell co-culture and tight junction formation.^9, 31^ The data shown in Figs. 2 and 3 validates that incorporating these components leads to a functional alveolar-capillary barrier. We then used this validated system to investigate the biomechanical mechanisms of VILI. First, we found that four hours of cyclic injurious stretch (20% radial strain, volutrauma) results in a significant loss of tight junction integrity as measured by reduced electrical impedance measurements (TEER; Fig. 4A). These results are consistent with previous studies which demonstrated that high cyclic stretch magnitudes result in a loss of barrier function as measured by albumin levels and TEER and decreased tight junction protein content^33-36^. In addition, Birukov et al. demonstrated that the deleterious effects of cyclic stretch are most severe within the first few minutes of cycling^35^ which is also consistent with our data where injurious levels of radial strain result in a significant decline in tight junction integrity within 30 minutes (Fig. 4A). After cessation of volutrauma, we observed a rapid increase in TEER measurements within 2 hours indicating that once injurious stretching is halted the barrier is capable of rapidly recovering. In support of this conclusion, previous studies have shown that following high tidal ventilation, lungs are able to reverse plasma membrane rupture, lung permeability and inflammation^37, 38^.

Atelectrauma is another well studied form of VILI and although previous studies investigated how atelectrauma alters cell death and substrate attachment^18, 25, 39^, our novel VOC monitors the effect of atelectrauma on transepithelial/endothelial barrier resistance in real-time during both injury and recovery. Both high and low frequency lung reopening resulted in rapid and significant drop in barrier function as measured by TEER (Fig. 5A) and barrier disruption was maintained during 2 hours of atelectrauma. In contrast to volutrauma, which begins to repair within one hour, the epithelium exposed to atelectrauma repairs more slowly. Specifically, TEER measurements after cessation of atelectrauma continue to be statistically lower than the zero-hour timepoints for up to 2 hours (Fig. 5B). This data suggests that atelectrauma induces a more severe injury that cannot recover as rapidly. Much work has been done by our group and others to investigate how cell matrix, temperature, cytoskeletal remodeling, and cell confluence impact cell injury during atelectrauma.^11, 12, 18, 24, 40, 41^ A common finding in these studies is that lower reopening velocities (lower frequency) lead to more plasma membrane disruption and cell death than atelectrauma at higher velocities/frequencies. Although we confirmed this inverse relationship between cell death and reopening velocity/frequency in this study (Fig. 5C), we did not observe a statistically significant difference in barrier disruption as measured by TEER between the high and low frequencies (Fig. 5B). Atelectrauma involves the generation of several complex mechanical forces and previous studies^24, 42^ have demonstrated that the large pressure gradients generated during slow airway reopening cause more plasma membrane disruption. However, the data in Fig 5B indicate these pressure gradients may not be the forces responsible for disrupting tight junctions. Future studies could therefore use computational technqiues^43^ to determine which mechanical force generated during airway reopening is responsible for barrier disruption.

While many investigators have studied how volutrauma or atelectrauma alone alter cell injury in-vitro^44^, to our knowledge Takayama et al.^45^ are the only group that used in-vitro microfluidics to evaluate cell injury during the simultaneous application of volutrauma and atelectrauma. Although the device used in that study utilized monocultures of transformed human or murine epithelial cells, a homogenous PDMS membrane, a short duration of injury (<5 min) and could only measure cell morphology/detachment, it did indicate that atelectrauma results in more cell detachment than volutrauma^45^. In this study, we developed a novel VOC that makes significant improvements in capturing important physiological and clinical aspects of VILI including the use of a deformable fibrous membrane that mimics the extracellular matrix, co-culture of primary human (non-transformed) epithelial and endothelial cells, real-time measurement of a clinically relevant parameter (barrier integrity) and application of injurious forces on clinically relevant time scales (i.e. hours to days). In addition, the VOC allows for important advantages as it can measure barrier function in real-time during both injury and recovery. Using the VOC, we demonstrated that concurrent application of volutrauma and atelectrauma for several hours results in barrier disruption and recovery patterns that are similar to atelectrauma alone (Fig 6A and 6B). Specifically, we observed a rapid and significant decline in barrier function during simultaneous volutrauma and atelectrauma as well as a slower recovery period. Interestingly, barrier disruption due to atelectrauma only and combined volutrauma and atelectrauma were not statistically different at any time point (Fig 6C and 6D). Therefore, our data strongly suggest that atelectrauma is driving mechanical forces responsible for injury/barrier disruption under simultaneous conditions and that atelectrauma is more injurious than volutrauma. This conclusion is consistent with previous work by Hussein et al. who found that ventilation of partially fluid filled lungs (atelectrauma) causes more injury than over-distending an air-filled lung (volutrauma)^41^. Furthermore, there is evidence that cyclic reopening (atelectrauma) and overdistention (volutrauma) work synergistically. In vivo, Seah et al. demonstrated that a combination of high tidal volume and no PEEP result in a dramatic decrease in the recruitablility of the lung as measured by lung stiffness^46^. More recently, a computational model suggests that the stresses of lung recruitment/decruitment are the primary instigating damage mechanism due to disruption of cell-cell and cell-substrate adhesions^47^ and that when coupled with overdistension, it leads to exacerbation of injury. This may have important therapeutic implications since previous studies have documented the cell mechanical changes needed to prevent atelectrauma^12, 43^.

Although our VOC model of the alveolar-capillary barrier measures how VILI impacts cell injury and recovery, our model does not replicate all aspects of in vivo lung physiology. First, we found that an air-liquid interface (ALI) culture was not necessary for tight junction formation and therefore used a submerged co-culture system. However, future studies could utilize ALI culture conditions if apical-basal transport was a desired endpoint. While we utilized a 20-30 *µ*m thick nanofiber mesh for ease of handling, we acknowledge that the basal lamina thickness in-vivo is ∼2-10 *µ*m. However, Fig. 3C demonstrates that our VOC forms a stronger barrier (i.e. higher TEER) under co-culture conditions as compared to monoculture conditions and therefore the membrane thickness does not inhibit cell-cell communication across the membrane. Finally, although the nanofiber mesh described here is stiffer than that of the human lung, it is still softer than other LOC scaffold fibers.^19^ Nevertheless, future studies could utilize a more biocompatible polymer (i.e. collagen) to lower the scaffold stiffness.

## Conclusion

In summary, we have developed a novel ventilator-on-a-chip (VOC) device that utilizes primary human epithelial and endothelial cells, a deformable nanofiber membrane, and microfluidics to simulate the complex mechanical forces that are responsible for lung injury during mechanical ventilation. We have used this system to demonstrate that atelectrauma is more injurious than volutrauma and that atelectrauma drives the injury response under simultaneous force application conditions. The VOC represents a powerful tool to study the mechanisms of ventilator induced lung injury and may also serve as a novel platform for drug discovery.

## Author Contributions (Contributor Roles Taxonomy, CRediT)

1. Conceptualization: BGZ, JAE, SNG
2. Data Curation: BGZ, JAE, SNG
3. Formal Analysis: BGZ, JAE, SNG
4. Funding Acquisition: JAE, SNG
5. Investigation: BGZ, VS
6. Methodology: BGZ, VS, CB, NHC, HP, JAE, SNG
7. Project Administration: JAE, SNG
8. Resources: NHC, HP, JAE, SNG
9. Software: BGZ, SNG
10. Supervision: JAE, SNG
11. Validation: BGZ, VS
12. Visualization: BGZ
13. Writing – Original Draft: BGZ, JAE, SNG
14. Writing – Review & Editing: BGZ, VS, CB, NHC, HP, JAE, SNG

## Acknowledgements

This work was supported by NIH Grant R56 HL142767 (J.A.E., S.N.G.), R01 HL142767 (J.AE. S.N.G.), and a Department of Defense Grant W81XWH-19-1-0210 (S.N.G., J.A.E.). We would like to acknowledge the contributions of the Center for Electron Microscopy and Analysis (CEMAS) at The Ohio State University for their technical assistance.

