## Supplemental Data for "A Micro-scale Humanized Ventilator-on-a-Chip to Examine the Injurious Effects of Mechanical Ventilation"

Supplementary Figures

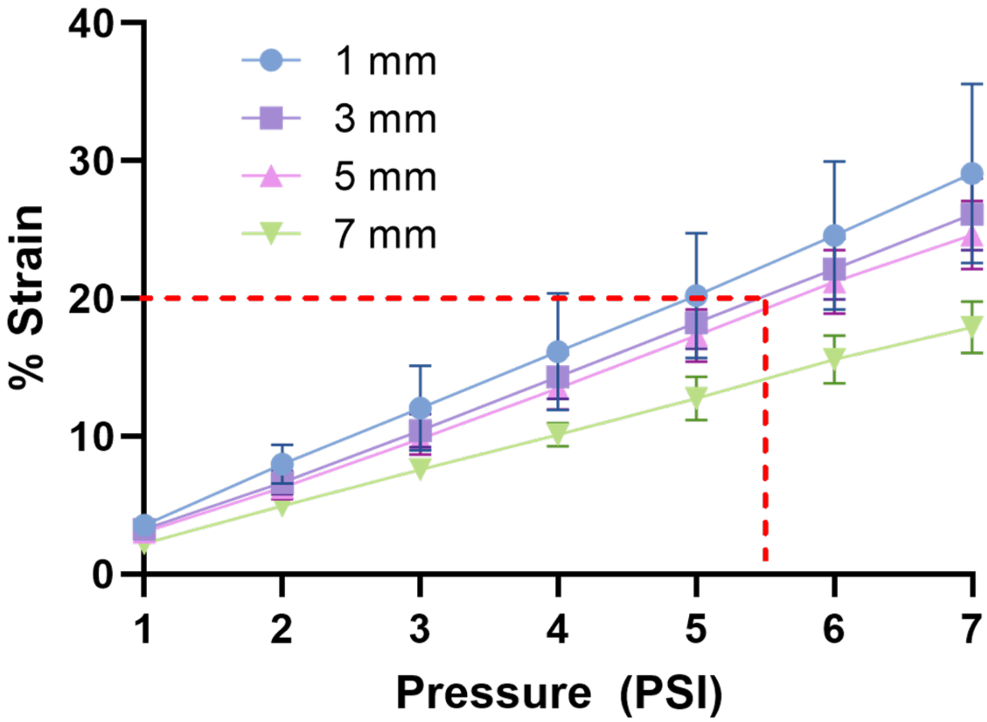


**Figure 1. VOC membrane stretch calibration.** Radial percent strain calculated by measuring the change in displacement of beads located 1, 3, 5, and 7 mm from the center of the membrane (Y-axis) vs. the corresponding vacuum pressure that linearly decreased from 0 to -7 PSI.
